# Genomic characterization of the non-O1/non-O139 *Vibrio cholerae* strain that caused a gastroenteritis outbreak in Santiago, Chile, 2018

**DOI:** 10.1101/835090

**Authors:** Mónica Arteaga, Juliana Velasco, Shelly Rodriguez, Maricel Vidal, Carolina Arellano, Francisco Silva, Leandro J. Carreño, Roberto Vidal, David A. Montero

**Affiliations:** Servicio de Urgencia Infantil, Hospital Clínico de la Universidad de Chile “Dr. José Joaquín Aguirre”, Santiago, Chile; Laboratorio de Salud Pública Ambiental y Laboral, Secretaría Regional Ministerial de Salud Región Metropolitana, Santiago, Chile; Programa de Microbiología y Micología, Instituto de Ciencias Biomédicas, Facultad de Medicina, Universidad de Chile, Santiago, Chile; Servicio de Laboratorio Clínico, Hospital Clínico de la Universidad de Chile “Dr. José Joaquín Aguirre”, Santiago, Chile; Programa de Inmunología, Instituto de Ciencias Biomédicas, Facultad de Medicina, Universidad de Chile, Santiago, Chile; Instituto Milenio de Inmunología e Inmunoterapia, Facultad de Medicina, Universidad de Chile, Santiago, Chile

**Keywords:** non-O1/non-O139*Vibrio cholerae*, gastroenteritis outbreak, type III secretion system (T3SS), type VI secretion system (T6SS), multidrug resistance genomic island

## Abstract

*Vibrio cholerae* is a human pathogen, which is transmitted by the consumption of contaminated food or water. *V. cholerae* strains belonging to the serogroups O1 and O139 can cause cholera outbreaks and epidemics, a severe life-threatening diarrheal disease. In contrast, serogroups other than O1 and O139, denominated as non-O1/non-O139, have been mainly associated with sporadic cases of moderate or mild diarrhea, bacteremia and wound infections. Here we investigated the virulence determinants and phylogenetic origin of a non-O1/non-O139 *V. cholerae* strain that caused a gastroenteritis outbreak in Santiago, Chile, 2018. We found that this outbreak strain lacks the classical virulence genes harboured by O1 and O139 strains, including the cholera toxin (CT) and the toxin-coregulated pilus (TCP). However, this strain carries genomic islands (GIs) encoding Type III and Type VI secretion systems (T3SS/T6SS) and antibiotic resistance genes. Moreover, we found these GIs are wide distributed among several lineages of non-O1/non-O139 strains. Our results suggest that the acquisition of these GIs may enhance the virulence of non-O1/non-O139 strains that lack the CT and TCP-encoding genes. Our results highlight the pathogenic potential of these *V. cholerae* strains.

**DATA SUMMARY:** Sequence data were submitted to GenBank under the accession number SRLP00000000. The authors confirm that all supporting data and protocols have been provided within the article or through supplementary data files.

**Data statement:** All supporting data, code and protocols have been provided within the article or through supplementary data files. Four supplementary tables are available with the online version of this article.

## INTRODUCTION

*V. cholerae* strains belonging to serogroups O1 and O139 are known to cause cholera outbreaks and epidemics. These serogroups generally produce the cholera toxin (CT) and the toxin coregulated pilus (TCP), which are responsible for secretory diarrhea and intestinal colonization, respectively [1]. Serogroups other than O1 and O139, called non-O1/non-O139, typically lack the CT and TCP-encoding genes [2]. However, several non-O1/non-O139 *V. cholerae* strains harbour additional virulence factors that contribute to pathogenicity [3]. In fact, non-O1/non-O139 *V. cholerae* strains have been isolated from sporadic cases of gastroenteritis, bacteremia and wound infections [1, 2, 4]. To date, the pathogenic mechanisms of non-O1/non-O139 *V. cholerae* strains have not been fully investigated.

There were several cholera outbreaks in South America in the 1990s [5]. In Chile, the last reported cases of cholera corresponded to the 1997–1998 outbreak of San Pedro de Atacama. Until 2017, only sporadic cases of gastroenteritis or bacteremia caused by non-O1/non-O139 *V*. cholerae strains had been reported in the country [6, 7]. On 2 August 2018, an outbreak of acute gastroenteritis started in Santiago, Chile. As of 11 March 2019, 70 gastroenteritis cases were reported, of which 13 required hospitalization, with diarrhea, nausea and vomiting being the main symptoms. As a result, an epidemiological study of the outbreak was performed by the National Reference Laboratory at the Public Health Institute of Chile (Instituto de Salud Pública de Chile, ISP). The official report of the outbreak indicated that among the patients, *V. cholerae* infection was confirmed by stool culture in 55/70 (78.5%), while FilmArray was positive for *V. cholerae* in 9/70 (12.8%) but without culture confirmation, and stool culture and FilmArray were negative for *V. cholerae* in 6/70 (8.5%). Additional analyses performed on the 45 *V. cholerae* strains isolated during the outbreak indicated that they are non-toxigenic and non-O1/non-O139. Importantly, a PFGE analysis showed that 42/45 strains were clonal (pulsotype 089), indicating that this clone was the main etiologic agent of the outbreak, although the source of contamination could not be determined [8]. Here, we performed comparative genomic and phylogenetic analyses to decipher the virulence determinants and phylogenetic origin of this outbreak strain.

## METHODS

### Antimicrobial drug susceptibility test

The disk diffusion method or broth dilution was performed according to Clinical and Laboratory Standards Institute guidelines [9].

### Genome sequencing

Genomic DNA of the *V. cholerae* str. Santiago-089 was extracted using the Wizard genomic DNA purification kit (Promega, USA) and sequenced at MicrobesNG (University of Birmingham, UK) using the Illumina MiSeq or HiSeq 2500 technology platforms with 2×250 bp paired-end reads. Draft genomes were provided after trimming low-quality ends and assembling reads with SPAdes 3.10 [10]. Assembly statistics were obtained with Quast v4.6.3 [11]. Contigs shorter that 200 nt were removed and sequences were deposited in GenBank under the accession number SRLP00000000.

### Publicly available genome sequences

A total of 69 genome sequences of *V. cholerae* strains were downloaded from GenBank on 1 June 2019. Genome accession numbers are listed in Table S1 (available in the online version of this article). Sequence management and blastn searches were performed using the Geneious software (v11.0.5; Biomatters).

### Phylogenetic analysis

A maximum likelihood phylogenetic tree based on core single nucleotide polymorphisms (SNPs) of 70 complete or draft genomic sequences of *V. cholerae* strains was built using CSI Phylogeny 1.4 [12]. This analysis was performed using the default input parameters and *V. cholerae* str. N16961 as the reference genome. As a result, 146534 SNPs were identified in 2483145 positions found in all analysed genomes. The output file in Newick format was downloaded and used to visualize the phylogenetic tree in the Interactive Tree of Life tool [13]. The population structure of the strains was determined with RhierBAPS [14] using the 146534 SNPs. For this, two depth levels and a maximum clustering size of 14 (default=number of isolates/5; 70/5=14) were specified. MLST sequence types were determined using the MLST 2.0 tool [15].

##### Impact Statement

*V. cholerae* remains a major public health problem in many Asian and African countries. In Chile, only sporadic or imported cases of gastroenteritis caused by *V. cholerae* have been reported in the last 20 years. However, in 2018, a clonal non-toxigenic and non-O1/non-O139 *V. cholerae* strain caused a gastroenteritis outbreak in the country. Typically, non-O1/non-O139 *V. cholerae* strains lack the cholera toxin; therefore, they must use additional virulence factors to cause severe disease. Consistent with this, we analysed this outbreak strain and found that it harbours genomic islands encoding T3SS, T6SS and antibiotic resistance genes that could promote its virulence. Moreover, our results show that non-O1/non-O139 *V. cholerae* is a heterogeneous group where these virulence factors are widespread among different clades. Knowledge of the acquisition of mobile genetic elements and the genetic diversity of these pathogens in order to understand their evolution and virulence potential is highly valuable. Since permanent surveillance of these pathogens is needed, we propose the use of T3SS and T6SS genes as molecular risk markers.

### Detection of virulence genes

Virulence genes analysed in this study and their GenBank accession numbers are listed in Table S2. The presence/ absence of virulence genes was determined using the blastn algorithm implemented in the Geneious software (v11.0.5; Biomatters). The absence of a gene was defined as an identity and/or gene coverage of less than 80 and 60%, respectively. The heat map showing the presence, absence and variation of the virulence genes was drawn using the gplots package [16] in R [17].

### Comparative genomic analysis and identification of genomic islands

Identification and characterization of DNA regions with features of genomic islands were performed using REPuter [18], ISfinder [19] and tRNAscan-SE [20]. ORFs were determined using the Geneious software (v11.0.5; Biomatters) and rast server [21]. The ORFs of the GI*Vch*-T6SS_Santiago-089_ and GI*Vch*-MDR_Santiago-089_ are listed in Tables S3 and S4, respectively. The comparison of the genetic structure of genomic islands was performed using EasyFig [22]. Additionally, the presence/ absence of the genomic islands VPI-I (GenBank accession: NC_002505.1, positions 873242–915211), VPI-II (GenBank accession: NC_002505.1, positions 1895692–1952861), VSP-I (GenBankaccession: NC_002505.1, positions175343–189380) and VSP-II (GenBank accession: NC_002505.1, positions 520634–550262) and Phage CTXφ (GenBank accession: NC_002505.1, positions 1550108–1574355) were determined using brig [23].

## RESULTS

The *V. cholerae* strain characterized in this study, which we named the Santiago-089 strain, was isolated from a 12-year-old boy hospitalized with bloody diarrhea and abdominal pain. This was one of the first gastroenteritis cases of the outbreak. Moreover, a PFGE analysis performed at the ISP showed that this and another 41 strains isolated during the outbreak had the same pulsotype with indistinguishable macrorestriction patterns, indicating that they are clonal (personal communication of the National Reference Laboratory at the Public Health Institute of Chile, ISP).

Initially, an antimicrobial drug susceptibility test showed that the Santiago-089 strain is resistant to trimethoprim-sulfamethoxazole, erythromycin and nalidixic acid (Table 1). Next, the phylogeny of this outbreak strain was investigated. For this, its genomic DNA was sequenced, and the draft genome deposited in GenBank under the accession number SRLP00000000. In addition, a set of genomes of *V. cholerae* strains isolated worldwide that are available in GenBank were included in the phylogenetic analysis (Table S1). As shown in the maximum likelihood phylogenetic tree (Fig. 1a), the strains were clustered into 12 lineages. While lineages 4 and 5 clustered the O1, O139 and O65 strains, the rest of the lineages clustered only non-O1/non-O139 strains. In particular, the Santiago-089 strain was clustered in lineage 2 along with strains isolated from India, Bangladesh and Haiti. Moreover, MLST sequence types were consistent with the topology of the tree.

**Table 1.**
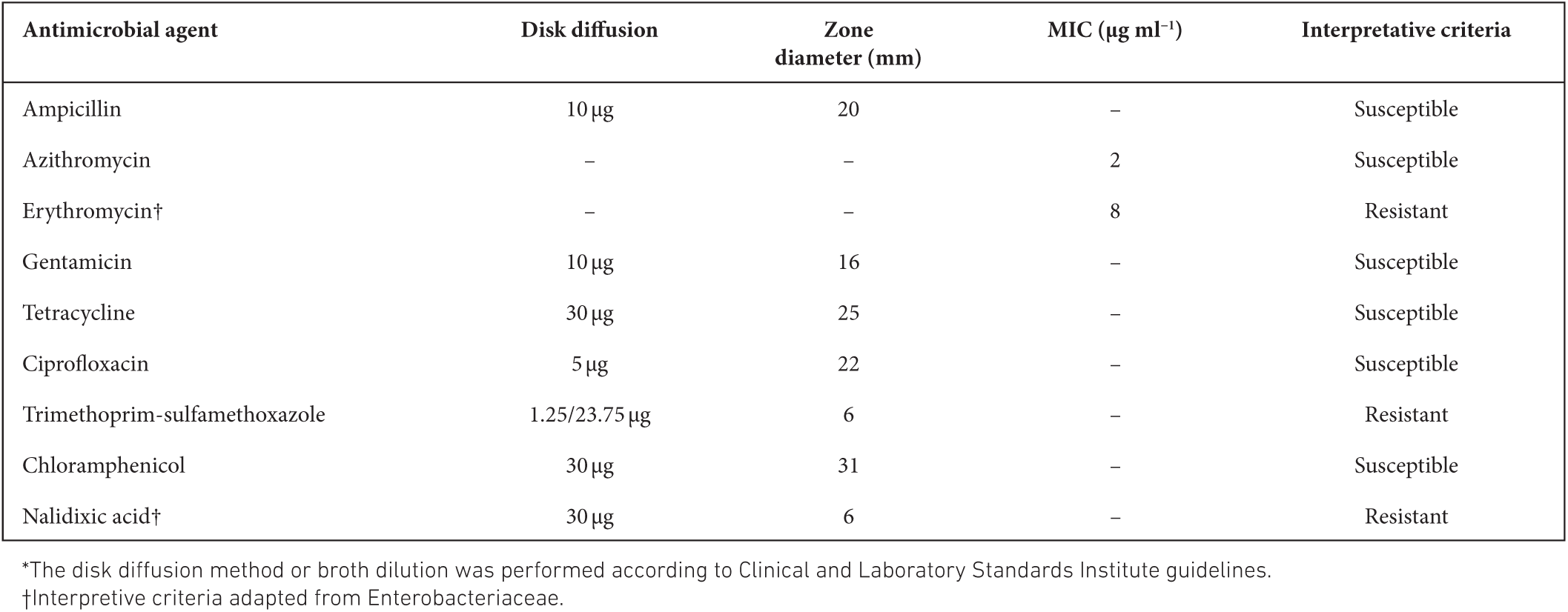
Antimicrobial resistance profile of the *V. cholerae* str. Santiago-089*

**Fig. 1.**
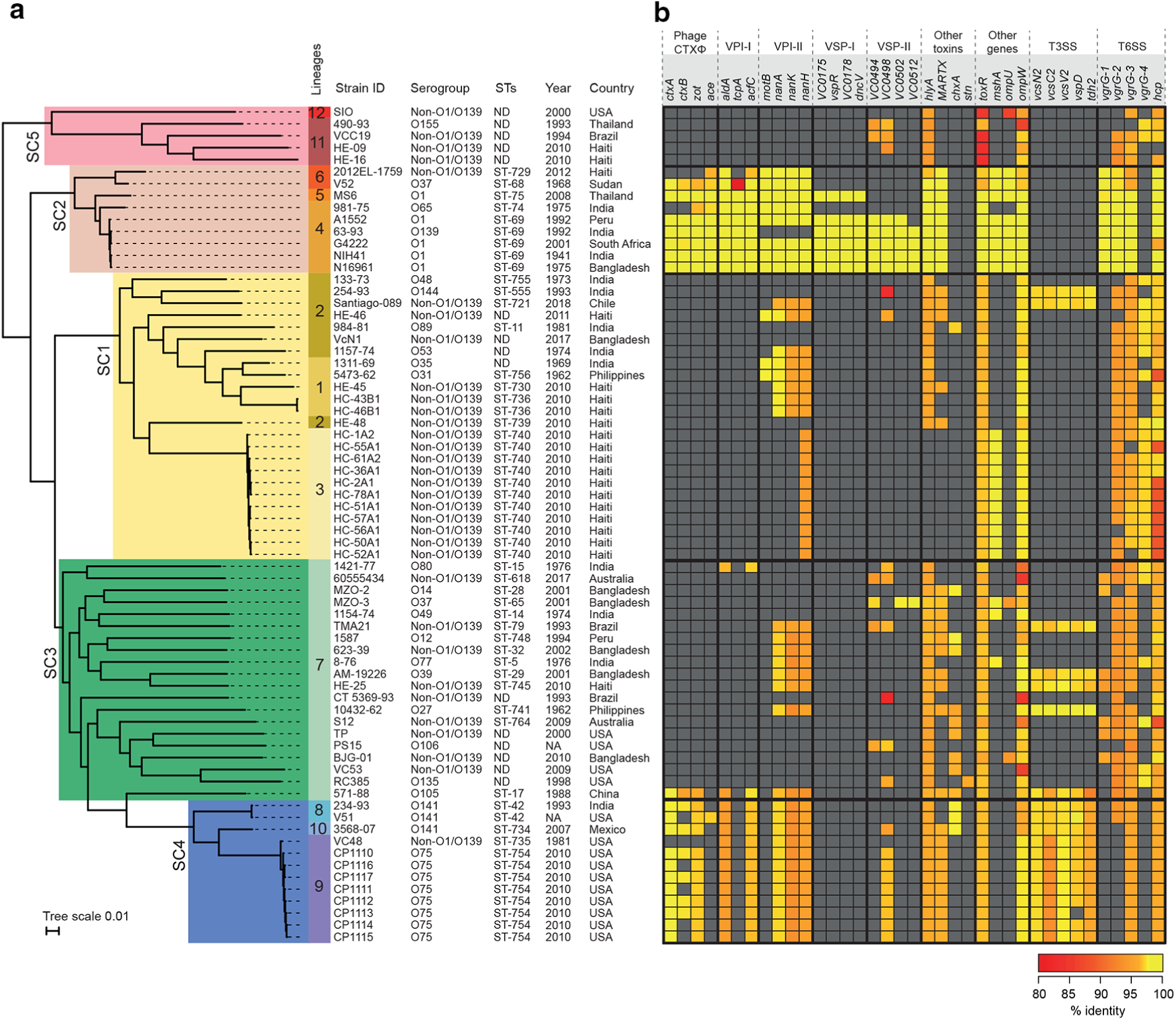
Phylogenetic relationship between 70 *V. cholerae* strains. (a) Maximum likelihood phylogenetic tree (midpoint rooted) based on whole-genome SNPs (146 534 SNPs within 2 483 145 positions, which were found in all analysed genomes). The genome of the *V. cholerae* str. N16961 was used as the reference. Bayesian analysis of population structure grouped the strains into five sequence clusters (SC; SC1 to SC5), which were further divided into 12 lineages. The epidemiological data of each strain is shown, including country and year of isolation. MLST sequence types (STs) are shown. (b) Heat map showing the presence, absence and variation of major virulence-associated genes distributed among the strains. Presence and variation (nucleotide identity levels, ranging from 80 to 100%) for each gene are indicated by colour intensity (red to yellow), as shown in the legend. The analysis was performed using blastn. Absence was defined as an identity and/or gene coverage of less than 80 and 60%, respectively and is indicated in grey.

This outbreak strain caused several hospitalizations but lacks the CT and, consequently other virulence genes must be contributing to its pathogenicity. Therefore, we analysed its genome searching for other virulence genes (Table S2). As expected, the Santiago-089 strain lacks the Phage CTXφ and the genomic islands VPI-I, VPI-II, VSP-I and VSP-II, which are generally carried by O1 and O139 strains (Fig. 2). However, this strain carries genes that encode toxins such HlyA and MARTX, and proteins of the Type III and Type VI secretion systems (T3SS/T6SS) (Fig. 1b). Similarly, most of the non-O1/non-O139 strains lack the Phage CTXφ and the GIs mentioned; rather, they carry genes encoding proteins of T3SS and T6SS. In contrast, O1/O139 strains harboured T6SS genes but not T3SS genes.

**Fig. 2.**
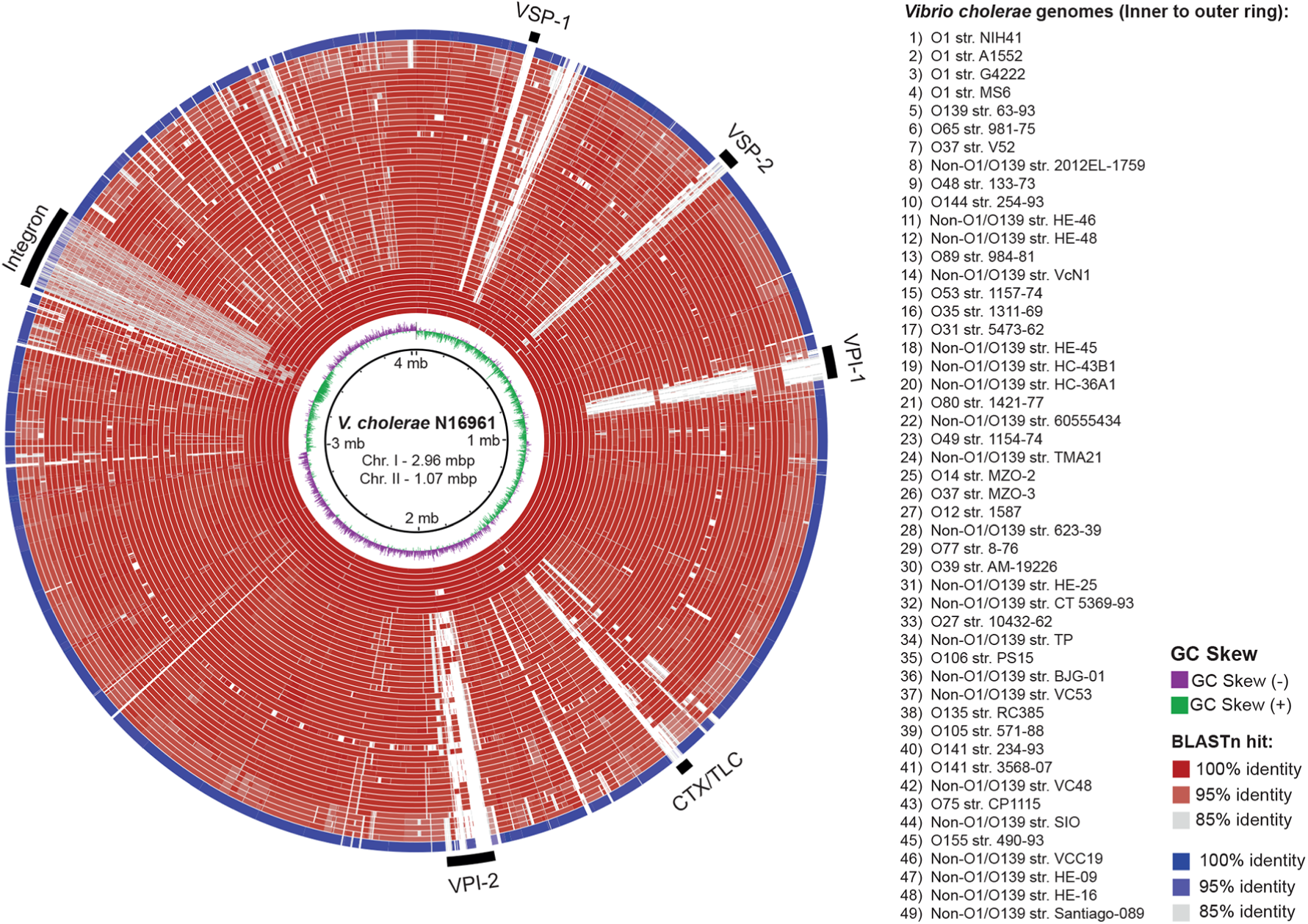
Comparison of the genomes of the *V. cholerae* str. Santiago-089 and 48 representative *V. cholerae* strains. The genomes are compared against the reference genome of *V. cholerae* str. N16961. The outermost ring (blue) shows the genome of *V. cholerae* str. Santiago-089. All other genomes are shown in red rings. blastn comparisons between reference genome and query genomes are shown as % identity according to the legend at the right. White regions indicate absence of genes or identity levels below 85%. The location of the major genomic islands is indicated as black bars. VSP, *Vibrio* seventh pandemic island; VPI, *Vibrio* pathogenicity island; CTX/TLC, cholera toxin/toxin-linked cryptic. The figure was prepared using brig [23].

We analysed the genetic context of the T3SS and T6SS genes in the Santiago-089 strain and found that they are located in GIs. The GI*Vch*-T3SS identified was previously reported in the *V. cholerae* str. AM-19226 and promotes colonization and infection [24]. We found that this GI of ∼64 kb is inserted in the tRNA-*ser* gene and located next to a nan-nag region involved in the sialic acid catabolism (Fig. 3a), which is also harboured by the VPI-II [25]. The second GI, which we named GI*Vch*-T6SS_Santiago-089_, was partially identified and contains a CRISPR-Cas region and genes that encode Hcp and VgrG alleles, which are structural T6SS components (Fig. 3b, Table S3). The GI*Vch*-T6SS_Santiago-089_ was also identified in the *V. cholerae* str. HC36A1. The Hcp from GI*Vch*T6SS_Santiago-089_ has 96.5% amino acid identity with the Hcp1 and Hcp2 alleles reported in *V. cholerae*. In contrast, the VgrG from GI*Vch*-T6SS_Santiago-089_ has 66.7, 66.6 and 59.4 % amino acid identity with the VgrG-1, VgrG-2 and VgrG-3 alleles reported in *V. cholerae*, respectively [26]. Therefore, we named this uncharacterized allele VgrG-4. Moreover, we found that Hcp and VgrG-4 alleles from GI*Vch*-T6SS_Santiago-089_ are also harboured by the GI*Vch*S12 [27]. Thus, the identification of T3SS genes and the VgrG-4 allele in non-O1/non-O139 strains of several lineages suggests these GIs are widely distributed (Fig. 1).

**Fig. 3.**
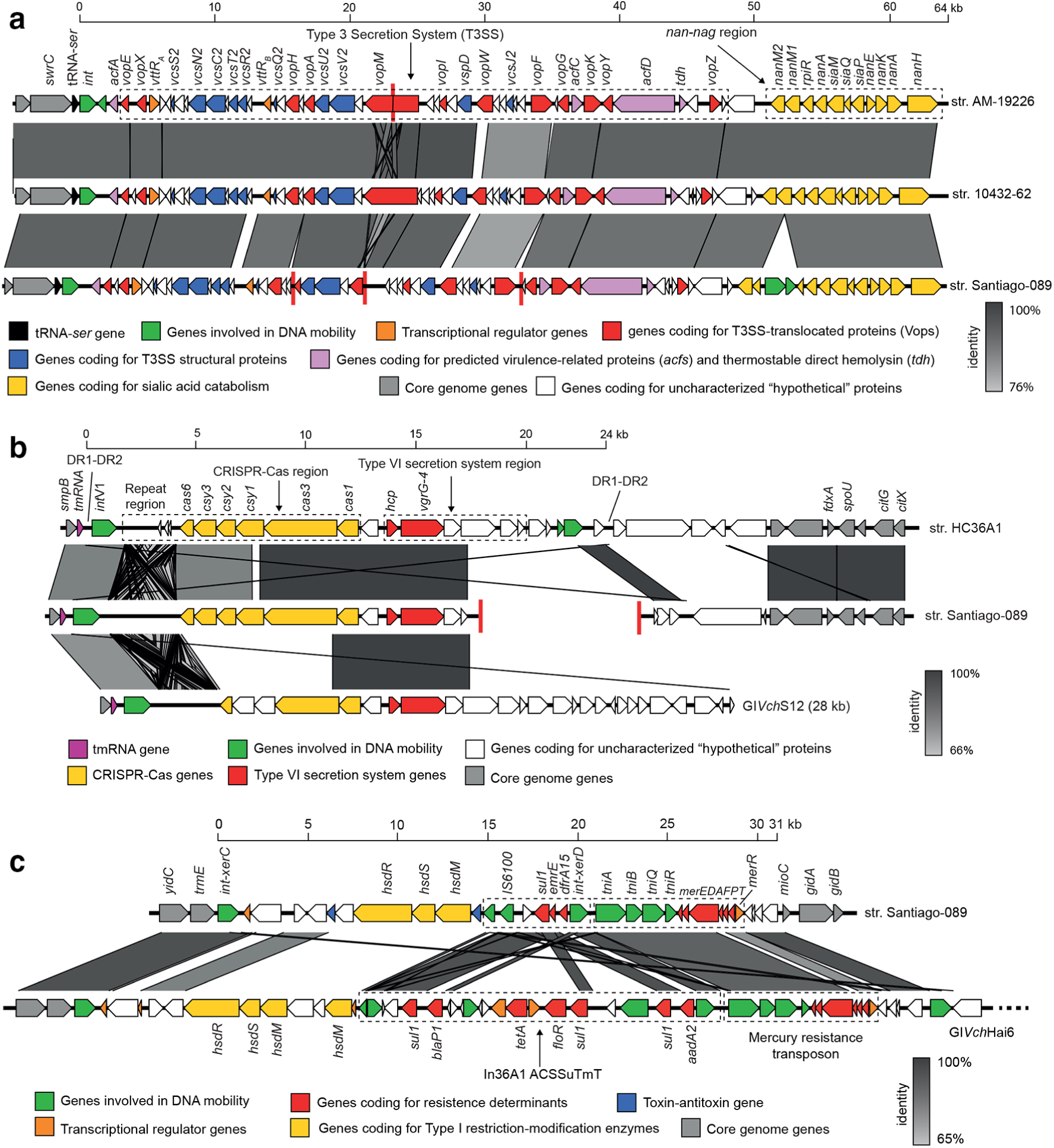
Comparison of the genetic structures of genomic islands carried by the *V. cholerae* str. Santiago-089. Predicted genes and the direction of transcription are represented as block arrows. Genes are colour-coded according to gene function, as indicated in the legends at the bottom. The names of some genes are indicated. Conserved regions are shaded in grey and the intensity of the colour indicates nucleotide identity levels, as indicated in the legends at the right. Contig boundaries are shown as red lines. (a) Genomic islands encoding a T3SS. GenBank accessions: GI*Vch*-T3SS_AM-19226_ (AATY02000004 and AATY02000003), GI*Vch*-T3SS_10432-62_ (CP010812), GI*Vch*-T3SS_Santiago-089_ (SRLP00000000). (b) Genomic islands encoding CRISPR-Cas and T6SS genes. GenBank accessions: GI*Vch*-T6SS_HC36A1_ (AXDR01000008.1), GI*Vch*-T6SS_Santiago-089_ (SRLP00000000), GI*Vch*S12 (KU722393). ORFs of the GI*Vch*-T6SS_Santiago-089_ are listed in Table S3. (c) Multidrug resistance (MDR) islands. The GI*Vch*-MDR_Santiago-089_ (GenBank accession: SRLP00000000) harbours the *sul1* and *dfrA15* genes, which confer resistance to sulfonamide and trimethoprim, respectively. ORFs of the GI*Vch*-MDR_Santiago-089_ are listed in Table S4. GI*Vch*Hai6 (GenBank accession: AXDR01000001).

Finally, since the Santiago-089 strain is multidrug-resistant (Table 1), we analysed its genome searching for genes or mutations that mediate its antimicrobial resistance profile. As a result, we identified a multidrug resistance (MDR) genomic island of ∼31 kb that we named GI*Vch*-MDR_Santiago-089_ (Fig. 3c, Table S4). This GI harbours the *sul1* and *dfrA15* genes, which confer resistance to sulfonamide and trimethoprim, respectively. The genetic structure of the GI*Vch*-MDR_Santiago-089_ is also relatively similar to the GI*Vch*Hai6 [28]. On the other hand, the Santiago-089 strain has the mutations S83I and A171S in the GyrA protein, which confer resistance to nalidixic acid [29] (Supplementary Material). Genes or mutations conferring resistance to erythromycin were not identified.

## DISCUSSION

*V. cholerae* remains a major public health problem, mainly in endemic areas of Asia and Africa, where access to adequate sanitation and safe drinking water is still limited [30]. In Chile, only sporadic or imported cases of gastroenteritis caused by non-O1/non-O139 *V. cholerae* strains have been reported in the last 20 years [6, 7]. Therefore, it was interesting to investigate the virulence determinants of the Santiago-089 strain.

The fact that a *V. cholerae* strain like the one characterized in the present study, which lacks the classical virulence factors (i.e. CT and CTP) yet causes a gastroenteritis outbreak with several hospitalized patients, represents a public health concern that justifies the permanent surveillance of these pathogens. Vaccination is a useful strategy to control cholera epidemics. However, licensed vaccines against *V. cholerae* target the O1 and O139 strains, and at this time there are no available vaccines against other serogroups [31]. This would be an important limitation to control a possible massive outbreak caused by non-O1/non-O139 strains. Moreover, the multidrug-resistant phenotype of the Santiago-089 strain, as well as the spread of MDR islands [28], are also epidemiological factors that need to be considered by public health authorities. In particular, the Santiago-089 strain was isolated from a case of bloody diarrhea. Although bloody diarrhea is not a common symptom caused by *V. cholerae*, there are some reports in which non-O1/non-O139 strains have caused it [32]. The above is of major importance for the diagnosis and treatment of these infections.

We show that the population structure of *V. cholerae* and particularly of the non-O1/non-O139 strains is heterogeneous, having a clear phylogenetic diversity and where known virulence factors are widely distributed among lineages. Indeed, we found that in the absence of the classic virulence factors, the non-O1/non-O139 strains have acquired several GIs encoding T3SS and T6SS, which may enhance their virulence. In fact, it has been suggested that the cumulative acquisition of pathogenicity islands may increase the virulence and contribute to the spread and emergence of some enteropathogens [33]. T3SS has a key role in the pathogenesis of *Vibrio parahaemolyticus* [34]. Although less studied, the role of T3SS in the pathogenesis of *V. cholerae* and other pathogenic *Vibrio* species has begun to be understood [24, 35]. For instance, the highly virulent non-O1/ non-O139 *V. cholerae* AM-19226 strain requires a functional T3SS for intestinal colonization in the infant mouse model [24, 36] and to cause fatal diarrhea in the infant rabbit model [37]. Similarly, the T6SSs are important virulence factors that promote the pathogenicity and environmental survival of *V. cholerae*. Current models of the T6SS structure suggest that a trimer of the VgrG protein is located at the tip of the apparatus that acts as a puncturing device against targeted cells [38]. As previously mentioned, three VgrG alleles have been reported in *V. cholerae:* VgrG-1, which exhibits actin cross-linking activity with cytotoxic effects on eukaryotic cells [39]; VgrG-2, which is essential for the anti-amoebae and anti-bacterial activities of the T6SS [26]; and VgrG-3, with anti-bacterial function by hydrolysing the cell wall of Gram-negative bacteria [40]. Moreover, we identified a new VgrG allele that is widely distributed among non-O1/ non-O139 strains. In future studies we will investigate the biological function of this new allele. Thus, T3SS and T6SS genes could be considered molecular risk markers for these pathogens and may be useful in epidemiological monitoring studies. In conclusion, this study highlights the pathogenic potential of the Santiago-089 strain as well as other non-O1/ non-O139 *V. cholerae* strains.

## Abbreviations

CT: cholera toxin;
GI: genomic island;
ISP: Public Health Institute of Chile;
MDR: multidrug resistance;
MLST: multilocus sequence typing;
ORF: open reading frame;
PFGE: pulsed-field gel electrophoresis;
SNP: single nucleotide polymorphisms;
TCP: toxin-coregulated pilus;
T3SS: type III secretion system;
T6SS: type VI secretion system.

## Acknowledgements

We wish to thank the National Reference Laboratory at the Public Health Institute of Chile for the epidemiological information of the Santiago-089 strain.

## Author contributions

The study was conceptualized by D.A. Montero and R. Vidal. Laboratory work was done by D.A. Montero, M. Arteaga, J. Velasco, S. Rodriguez, C. Arellano, M. Vidal and F. Silva. Genomic analysis was carried out by D.A. Montero, M. Arteaga, J. Velasco and L.J. Carreño. *V. cholerae* str. Santiago-089 was provided by F. Silva. The manuscript was drafted by D.A. Montero, M. Arteaga and J. Velasco and reviewed and edited by D.A. Montero, M. Vidal, L.J. Carreño and R. Vidal. All authors have read and approved the final version of the manuscript.

## Ethical statement

The microbiological study of the *V. cholerae* Santiago-089 strain was approved by written consent of the patient’s parents and the Ethics Committee of the Clinical Hospital University of Chile.

## Data Bibliography

1. The accession numbers of publicly available genomes of *V. cholerae* used in this study are summarized in Table S1 (2019).
2. The accession numbers of virulence genes analysed in this study are summarized in Table S2 (2019).

### Five reasons to publish your next article with a Microbiology Society journal

1. The Microbiology Society is a not-for-profit organization.
2. We offer fast and rigorous peer review – average time to first decision is 4–6 weeks.
3. Our journals have a global readership with subscriptions held in research institutions around the world.
4. 80% of our authors rate our submission process as ‘excellent’ or ‘very good’.
5. Your article will be published on an interactive journal platform with advanced metrics.

**Find out more and submit your article at microbiologyresearch.org.**

## References

1. Dutta D, Chowdhury G, Pazhani GP, Guin S, Dutta S et al. *Vibrio cholerae* non-O1, non-O139 serogroups and cholera-like diarrhea, Kolkata, India. Emerg Infect Dis 2013;19:464–467.

2. Octavia S, Salim A, Kurniawan J, Lam C, Leung Q et al. Population structure and evolution of non-O1/non-O139 Vibrio cholerae by multilocus sequence typing. PLoS One 2013;8:e65342.

3. Chatterjee S, Ghosh K, Raychoudhuri A, Chowdhury G, Bhattacharya MK et al. Incidence, virulence factors, and clonality among clinical strains of non-O1, non-O139 Vibrio cholerae isolates from hospitalized diarrheal patients in Kolkata, India. J Clin Microbiol 2009;47:1087–1095.

4. Deshayes S, Daurel C, Cattoir V, Parienti J-J, Quilici M-L et al. non-O1, non-O139 Vibrio cholerae bacteraemia: case report and literature review. Springerplus 2015;4:575.

5. Domman D, Quilici M-L, Dorman MJ, Njamkepo E, Mutreja A et al. Integrated view of *Vibrio cholerae* in the Americas. Science 2017;358:789–793.

6. Olivares F, Domínguez I, Dabanch J, Porte L, Ulloa MT et al. Bacteriemia POR Vibrio cholerae no-01/no-0139 que porta una región homóloga a la isla de patogenicidad VpaI-7. Rev Chil infectología 2019;36:392–395.

7. Montero D, Vidal M, Pardo M, Torres A, Kruger E et al. Characterization of enterotoxigenic Escherichia coli strains isolated from the massive multi-pathogen gastroenteritis outbreak in the Antofagasta region following the Chilean earthquake, 2010. Infect Genet Evol 2017;52:26–29.

8. Ministerio de Salud de Chile. Departamento de Epidemiología. Minuta: Situación epidemiológica de brote de diarrea aguda por Vibrio cholerae no toxigénico. Chile, año 2018-2019 2019 (Not published).

9. Clinical and Laboratory Standards Institute - CLSI. Methods for Antimicrobial Dilution and Disk Susceptibility Testing of Infrequently Isolated or Fastidious Bacteria. M45, 3rd ed. Wayne, PA: Clinical and Laboratory Standards Institute; 2015.

10. Bankevich A, Nurk S, Antipov D, Gurevich AA, Dvorkin M et al. SPAdes: a new genome assembly algorithm and its applications to single-cell sequencing. J Comput Biol 2012;19:455–477.

11. Gurevich A, Saveliev V, Vyahhi N, Tesler G. QUAST: quality assessment tool for genome assemblies. Bioinformatics 2013;29:1072–1075.

12. Kaas RS, Leekitcharoenphon P, Aarestrup FM, Lund O. Solving the problem of comparing whole bacterial genomes across different sequencing platforms. PLoS One 2014;9:e104984.

13. Letunic I, Bork P. Interactive tree of life (iTOL) V3: an online tool for the display and annotation of phylogenetic and other trees. Nucleic Acids Res 2016;44:W242–W245.

14. Tonkin-Hill G, Lees JA, Bentley SD, Frost SDW, Corander J. Rhier-BAPS: an R implementation of the population clustering algorithm hierBAPS. Wellcome Open Res 2018;3:93.

15. Larsen MV, Cosentino S, Rasmussen S, Friis C, Hasman H et al. Multilocus sequence typing of total-genome-sequenced bacteria. J Clin Microbiol 2012;50:1355–1361.

16. Warnes GR, Bolker B, Bonebakker L, Gentleman R, Huber W. R Package ‘gplots’ 2016.

17. R Core Team. R: A Language and Environment for Statistical Computing. Vienna, Austria: R Foundation for Statistical Computing; 2014.

18. Kurtz S et al. REPuter: the manifold applications of repeat analysis on a genomic scale. Nucleic Acids Res 2001;29:4633–4642.

19. Siguier P, Perochon J, Lestrade L, Mahillon J, Chandler M. ISfinder: the reference centre for bacterial insertion sequences. Nucleic Acids Res 2006;34:D32–D36.

20. Lowe TM, Eddy SR. TRNAscan-SE: a program for improved detection of transfer RNA genes in genomic sequence. Nucleic Acids Res 1997;25:955–964.

21. Aziz RK, Bartels D, Best AA, DeJongh M, Disz T et al. The RAST server: rapid annotations using subsystems technology. BMC Genomics 2008;9:75–15.

22. Sullivan MJ, Petty NK, Beatson SA. Easyfig: a genome comparison visualizer. Bioinformatics 2011;27:1009–1010.

23. Alikhan N-F, Petty NK, Ben Zakour NL, Beatson SA. Blast ring image generator (BRIG): simple prokaryote genome comparisons. BMC Genomics 2011;12:402.

24. Chaand M, Miller KA, Sofia MK, Schlesener C, Weaver JWA et al. Type three secretion system island-encoded proteins required for colonization by non-O1/non-O139 serogroup Vibrio cholerae. Infect Immun 2015;83:2862–2869.

25. Almagro-Moreno S, Boyd EF. Sialic acid catabolism confers a competitive advantage to pathogenic Vibrio cholerae in the mouse intestine. Infect Immun 2009;77:3807–3816.

26. Zheng J, Ho B, Mekalanos JJ. Genetic analysis of Anti-Amoebae and anti-bacterial activities of the type VI secretion system in Vibrio cholerae. PLoS One 2011;6:e23876.

27. Labbate M, Orata FD, Petty NK, Jayatilleke ND, King WL et al. A genomic island in Vibrio cholerae with VPI-1 site-specific recombination characteristics contains CRISPR-Cas and type VI secretion modules. Sci Rep 2016;6:36891.

28. Carraro N, Rivard N, Ceccarelli D, Colwell RR, Burrus V. IncA/C Conjugative plasmids mobilize a new family of multidrug resistance islands in clinical *Vibrio cholerae* non-O1/non-O139 isolates from Haiti. MBio 2016;7.

29. Zhou Y, Yu L, Li J, Zhang L, Tong Y et al. Accumulation of mutations in DNA gyrase and topoisomerase IV genes contributes to fluoroquinolone resistance in Vibrio cholerae 0139 strains. Int J Antimicrob Agents 2013;42:72–75.

30. Deen J, Mengel MA, Clemens JD. Epidemiology of cholera. Vaccine 2019.

31. O’Ryan M, Vidal R, del Canto F, Salazar JC, Montero D. Vaccines for viral and bacterial pathogens causing acute gastroenteritis: Part I: Overview, vaccines for enteric viruses and *Vibrio cholerae*. Hum Vaccin Immunother 2015;11:584–600.

32. Fraga SG, De Trejo AV, Pichel M, Figueroa S, Merletti G et al. Caracterización de aislamientos de Vibrio cholerae no-01, no-0139 asociados a cuadros de diarrea. Rev Argent Microbiol 2009;41:11–19.

33. Montero DA, Canto FD, Velasco J, Colello R, Padola NL et al. Cumulative acquisition of pathogenicity islands has shaped virulence potential and contributed to the emergence of LEE-negative Shiga toxin-producing *Escherichia coli* strains. Emerg Microbes Infect 2019;8:486–502.

34. Broberg CA, Calder TJ, Orth K. Vibrio parahaemolyticus cell biology and pathogenicity determinants. Microbes Infect 2011;13:992–1001.

35. Zhao Z, Liu J, Deng Y, Huang W, Ren C et al. The *Vibrio alginolyticus* T3SS effectors, Val1686 and Val1680, induce cell rounding, apoptosis and lysis of fish epithelial cells. Virulence 2018;9:318–330.

36. Tam VC, Serruto D, Dziejman M, Brieher W, Mekalanos JJ. A type III secretion system in Vibrio cholerae translocates a Formin/Spire Hybrid-like actin nucleator to promote intestinal colonization. Cell Host Microbe 2007;1:95–107.

37. Shin OS, Tam VC, Suzuki M, Ritchie JM, Bronson RT et al. Type III secretion is essential for the rapidly fatal diarrheal disease caused by non-O1, non-O139 Vibrio cholerae. MBio 2011;2:1–11.

38. Cascales E, Cambillau C. Structural biology of type VI secretion systems. Phil Trans R Soc B 2012;367:1102–1111.

39. Pukatzki S, Ma AT, Revel AT, Sturtevant D, Mekalanos JJ. Type VI secretion system translocates a phage tail spike-like protein into target cells where it cross-links actin. Proc Natl Acad Sci U S A 2007;104:15508–15513.

40. Brooks TM, Unterweger D, Bachmann V, Kostiuk B, Pukatzki S. Lytic Activity of the *Vibrio cholerae* Type VI Secretion Toxin VgrG-3 Is Inhibited by the Antitoxin TsaB. J Biol Chem 2013;288:7618–7625.

