## Supplementary information for "Genomic characterization of the non-O1/non-O139 *Vibrio cholerae* strain that caused a gastroenteritis outbreak in Santiago, Chile, 2018" (from another version of this preprint: version 3, 2019-11-11)

**Supplementary Table 1.** *Vibrio cholerae* genomes analyzed in this study

| Strain ID | Country | Origin / Host | Diagnostic | No. of contigs | GenBank Accession |
|---|---|---|---|---|---|
| 5473-62 | Philippines | Human | Diarrhea | 94 | JIDI00000000 |
| 1311-69 | India | Human | Diarrhea | 130 | JIDJ00000000 |
| 1157-74 | India | Human | Diarrhea | 151 | JIDL00000000 |
| HE-45 | Haiti | Environment | - | 29 | ALED00000000 |
| HC-43B1 | Haiti | Human | - | 20 | ALDP00000000 |
| HC-46B1 | Haiti | Human | - | 24 | AJSL00000000 |
| VcN1 | Bangladesh | Environment | - | 178 | PDNJ00000000 |
| 133-73 | India | Human | Diarrhea | 118 | JIDK00000000 |
| 984-81 | India | Human | Diarrhea | 117 | JMBM00000000 |
| 254-93 | India | Human | Diarrhea | 114 | JMBP00000000 |
| HE-46 | Haiti | Environment | - | 143 | MIPM00000000 |
| HE-48 | Haiti | Environment | - | 18 | AFOR00000000 |
| HC-1A2 | Haiti | Human | - | 19 | AJRO00000000 |
| HC-2A1 | Haiti | Human | - | 206 | AFOT00000000 |
| HC-36A1 | Haiti | Human | - | 17 | AXDR00000000 |
| HC-61A2 | Haiti | Human | - | 24 | AJRU00000000 |
| HC-78A1 | Haiti | Human | - | 172 | AGUZ00000000 |
| HC-50A1 | Haiti | Human | - | 254 | ALDS00000000 |
| HC-51A1 | Haiti | Human | - | 249 | ALDT00000000 |
| HC-52A1 | Haiti | Human | - | 271 | ALDU00000000 |
| HC-55A1 | Haiti | Human | - | 243 | ALDV00000000 |
| HC-56A1 | Haiti | Human | - | 217 | ALDW00000000 |
| HC-57A1 | Haiti | Human | - | 237 | ALDY00000000 |
| G4222 | South Africa | Human | Cholera | 21 | ANNB01000000 |
| NIH41 | India | Human | NA | 111 | JIDH00000000 |
| 981-75 | India | Human | Diarrhea | 152 | JIDM00000000 |
| 63-93 | India | Human | Diarrhea | 81 | JMBN00000000 |
| A1552 | Peru | Human | Cholerae | 2 | CP025936, CP025937 |
| N16961 | Bangladesh | Human | Cholerae | 2 | NZ_CP028827.1, NZ_CP028828.1 |
| MS6 | Thailand | Human | Diarrhea | 2 | AP014524, AP014525 |
| 2012EL-1759 | Haiti | Environment | - | 68 | JNEW00000000 |
| V52 | Sudan | Human | - | 111 | AAKJ00000000 |
| 1154-74 | India | Human | Diarrhea | 1 | CP010811 |
| 10432-62 | Philippines | Human | Diarrhea | 1 | CP010812 |
| PS15 | USA | Environment | - | 131 | AIJR00000000 |
| 8_76 | India | Human | Diarrhea | 82 | JIDN00000000 |
| 1421-77 | India | Human | Diarrhea | 118 | JMBL00000000 |
| 571-88 | China | Human | Diarrhea | 117 | JIDO00000000 |
| 60555434 | Australia | Human | Dysuria | 95 | QKWT00000000 |
| MZO-2 | Bangladesh | Human | - | 157 | MIKJ00000000 |
| TP | USA | Environment | - | 291 | MIPK00000000 |
| VC53 | USA | Oyster | - | 247 | MIOU00000000 |
| 1587 | Peru | Human | - | 114 | AAUR00000000 |
| 623-39 | Bangladesh | Environment | - | 80 | AAWG00000000 |
| AM-19226 | Bangladesh | Human | - | 62 | AATY00000000 |
| CT 5369-93 | Brazil | Environment | - | 269 | ADAL00000000 |
| MZO-3 | Bangladesh | Human | - | 77 | AAUU00000000 |
| RC385 | USA | Environment | - | 97 | AAKH00000000 |
| TMA21 | Brazil | Environment | - | 20 | ACHY00000000 |
| HE-25 | Haiti | Environment | - | 13 | ALEC00000000 |
| BJG-01 | USA | NA | - | 311 | AFOU00000000 |
| 234-93 | India | Human | Diarrhea | 129 | JMBO00000000 |
| V51 | USA | Human | - | 143 | AAKI00000000 |
| CP1110 | USA | Human | Cholera | 184 | AMWF00000000 |
| CP1111 | USA | Human | Cholera | 199 | AMWS00000000 |
| CP1112 | USA | Human | Cholera | 193 | AMWT00000000 |
| CP1113 | USA | Human | Cholera | 186 | AMWU00000000 |
| CP1114 | USA | Human | Cholera | 199 | AMWV00000000 |
| CP1115 | USA | Human | Cholera | 183 | AMWR00000000 |
| CP1116 | USA | Human | Cholera | 207 | ANNM00000000 |
| CP1117 | USA | Human | Cholera | 183 | AMWW00000000 |
| VC48 | USA | Oyster | - | 152 | MIOT00000000 |
| 3568-07 | Mexico | Fresh Cheese | - | 142 | MIPL00000000 |
| VCC19 | Brazil | Environment | - | 54 | ATEV00000000 |
| 490-93 | Thailand | Human | Diarrhea | 274 | JIDQ00000000 |
| HE-09 | Haiti | Environment | - | 258 | AFOP00000000 |
| HE-16 | Haiti | Environment | - | 326 | ALEB00000000 |
| SIO | USA | Environment | - | 150 | MIPJ00000000 |
| S12 | Australia | Environment | - | 83 | MDST01000000 |

**Supplementary Table 2.** Virulence associated genes analyzed in this study

| Virulence genes | Gene name | GenBank accession | Nucleotide positions |
|---|---|---|---|
| **Filamentous phage CTXΦ** | | | |
| *ctxA* | Cholera enterotoxin, A subunit | LT906614.1 | c1568126-1567350 |
| *ctxB* | Cholera enterotoxin, B subunit | LT906614.1 | c1567353-1566979 |
| *zot* | Zonular occludens toxin | LT906614.1 | c1569424-1568225 |
| *ace* | Accessory cholera enterotoxin | LT906614.1 | c1569702-1569409 |
| **Vibrio pathogenicity island-I (VPI-I)** | | | |
| *aldA* | Aldehyde dehydrogenase | LT906614.1 | 875278-876903 |
| *tcpA* | Toxin-coregulated pilin | AF452571.1 | |
| *acfC* | Accessory colonization factor AcfC | LT906614.1 | 903385-904155 |
| **Vibrio pathogenicity island-II (VPI-II)** | | | |
| *motB-like* | Chemotaxis protein MotB-related protein | LT906614.1 | c1903654-1902920 |
| *nanA* | N-acetylneuraminate lyase | LT906614.1 | c1927114-1926218 |
| *nanK* | N-acetylmannosamine kinase | LT906614.1 | 1930905-1931768 |
| *nanH* | Neuraminidase | LT906614.1 | 1933253-1935676 |
| **Vibrio Seventh Pandemic Island-I (VSP-I)** | | | |
| *VC0175* | Deoxycytidylate deaminase-related protein | LT906614.1 | c176941-175343 |
| *vspR* | *V. cholerae* seventh pandemic regulator | LT906614.1 | 177861-178424 |
| *VC0178* | Patatin-related protein | LT906614.1 | 179338-180405 |
| *dncV* | Di-nucleotide cyclase Vibrio | LT906614.1 | 180419-181729 |
| **Vibrio Seventh Pandemic Island-II (VSP-II)** | | | |
| *VC0494* | Hypothetical protein | LT906614.1 | 528305-528949 |
| *VC0498* | Ribonuclease HI | LT906614.1 | 530684-531124 |
| *VC0502* | Type IV pilin | LT906614.1 | c534722-534198 |
| *VC0512* | Methyl-accepting chemotaxis protein | LT906614.1 | 541318-542907 |
| **Other toxins** | | | |
| *hlyA* | Vibrio cholerae cytolysin | LT906615.1 | 237585-239810 |
| *rtxA* | MARTX (multifunctional autoprocessing RTX toxin) | AE003852.1 | 1550108-1563784 |
| *chxA* | Cholix toxin | AB754476.1 | |
| *stn* | Heat-stable enterotoxin, NAG-ST | M85198.1 | |
| **Other virulence associated genes** | | | |
| *toxR* | ToxR, regulator of the expression of virulence factors | LT906614.1 | c1047905-1047185 |
| *mshA* | Mannose-sensitive hemagglutinin (MSHA), a type 4 pilus | LT906614.1 | 435450-435986 |
| *ompU* | Outer membrane protein U | KF434513.1 | |
| *ompW* | Outer membrane protein W | LT906615.1 | c820096-819681 |
| **Type III secretion system (T3SS)** | | | |
| *vcsN2* | Type III secretion cytoplasmic ATPase | CP010812.1 | c2624653-2623391 |
| *vcsC2* | Type II/III secretion system protein | CP010812.1 | c2626110-2624653 |
| *vcsV2* | Type III secretion protein | CP010812.1 | c2635653-2633770 |
| *vspD* | Putative secreted protein D | CP010812.1 | c2644010-2642961 |
| *tdh2* | Thermostable direct hemolysin 2 | CP010812.1 | 2659186-2659728 |
| **Type VI secretion system (T6SS)** | | | |
| *vgrG-1* | Valine-glycine repeat protein G 1 | LT906614.1 | 1510289-1513780 |
| *vgrG-2* | Valine-glycine repeat protein G 2 | LT906615.1 | 20426-22510 |
| *vgrG-3* | Valine-glycine repeat protein G 3 | LT906615.1 | 137579-140632 |
| *vgrG-4* | Valine-glycine repeat protein G 4 | KU722393.1 | 13902-15878 |
| *hcp* | Hemolysin coregulated protein | LT906614.1 | 1509643-1510161 |

**Supplementary Table 3.** Open reading frames localized within GI*Vch*-T6SS$_{\text{Santiago-089}}$ (GenBank accession: SRLP00000000)

| ORF# | Position (bp) * | Denomination in transposable element | Closest informative protein match | Identities | Accession No. of homologue | Function to closest related protein. Comments |
|---|---|---|---|---|---|---|
| | 1 - 367 | *ssrA* tmRNA | | | | |
| 1 | 534 – 1,772 | Integrase | Tyrosine-type recombinase/integrase | 411/412 (99%) | WP_032480643.1 | DNA mobility |
| | c2,129 – 5,276 | CRISPR with 53 repeat units repeat region | | | | |
| 2 | c5,432 – 6,070 | *cas6* | Type I-F CRISPR-associated endoribonuclease Cas6/Csy4 | 211/212 (99%) | WP_142735065.1 | CRISPR-Cas system |
| 3 | c6,080 – 7,111 | *csy3* | Type I-F CRISPR-associated protein Csy3 | 343/343 (100%) | WP_142576943.1 | CRISPR-Cas system |
| 4 | c7,123 – 8,010 | *csy2* | Type I-F CRISPR-associated protein Csy2 | 295/295 (100%) | WP_142735064.1 | CRISPR-Cas system |
| 5 | c8,007 – 9,284 | *csy1* | Type I-F CRISPR-associated protein Csy1 | 425/425 (100%) | WP_142735063.1 | CRISPR-Cas system |
| 6 | c9,284 – 12,652 | *cas3* | Type I-F CRISPR-associated helicase Cas3 | 1120/1122 (99%) | WP_071192023.1 | CRISPR-Cas system |
| 7 | c12,649 – 13,638 | *cas1* | Type I-F CRISPR-associated endonuclease Cas1 | 327/329 (99%) | WP_119788720.1 | CRISPR-Cas system |
| 8 | c13,753 – 14,547 | Hypothetical protein | WYL domain-containing protein | 264/264 (100%) | WP_000880612.1 | Predicted DNA-binding transcriptional regulator contains HTH and WYL domains |
| 9 | 14,963 – 15,481 | *hcp* | Hcp family type VI secretion system effector | 172/172 (100%) | WP_001142919.1 | Type VI secretion system effector |
| 10 | 15,608 – 17,584 | *vgrG-4* | VgrG-2 protein | 428/620 (69%) | WP_000212125.1 | Type VI secretion system effector |
| 11 | 17,584 – 18,372 | Hypothetical protein | DUF4123 domain-containing protein | 260/262(99%) | WP_033932339.1 | Domain of unknown function (DUF4123) |
| 12 | 18,362 – 18,670 | Hypothetical protein | Hypothetical protein | 98/98 (100%) | WP_142735597.1 | Unknown |

* c: indicates ORFs transcribed on the complementary strand.

**Supplementary Table 4.** Open reading frames localized within GI*Vch*-MDR$_{Santiago-089}$ (GenBank accession: SRLP00000000)

| ORF# | Position (bp) * | Denomination in transposable element | Closest informative protein match | Identities | Accession No. of homologue | Function to closest related protein. Comments |
|---|---|---|---|---|---|---|
| | trmE gene | | | | | |
| 1 | 1 – 1,170 | Int-xerC | Site-specific integrase | 385/389 (99%) | WP_000703233.1 | DNA mobility |
| 2 | c1,414 – 1,758 | Hypothetical protein | Helix-turn-helix domain-containing protein | 114/114 (100%) | WP_063108740.1 | Predicted transcriptional regulator |
| 3 | c1,818 – 3,488 | Hypothetical protein | DUF927 domain-containing protein | 555/556 (99%) | WP_095482111.1 | Domain of unknown function (DUF927) |
| 4 | 4,226 – 4,903 | Hypothetical protein | Hypothetical protein | 224/224 (100%) | WP_032478679.1 | Unknown |
| 5 | c4,961 – 5,995 | Hypothetical protein | Virulence RhuM family protein | 337/344 (98%) | WP_023135447.1 | Unknown |
| 6 | c6,006 – 6,476 | Toxin-Antitoxin gene | Type II toxin-antitoxin system death-on-curing family toxin | 156/156 (100%) | WP_023135448.1 | Toxin-antitoxin system |
| DR | 6,228- 6,242<br>28,579 – 28,593 | AGC--GCCAGTGAAGAGCG<br>AGCCCGCCAGTGAAGAGCG | | | | |
| 7 | c6,478 – 7,491 | Hypothetical protein | DNA-binding protein | 337/337 (100%) | EBV0095603.1 | Unknown |
| 8 | c7,484 – 10,729 | *hsdR* | Type I restriction endonuclease subunit R | 1081/1081 (100%) | EBV0095602.1 | Type I restriction-modification enzymes |
| 9 | c10,729 – 12,018 | *hsdS* | Restriction endonuclease subunit S | 429/429 (100%) | EBV0095601.1 | Type I restriction-modification enzymes |
| 10 | c12,015 – 13,997 | *hsdM* | SAM-dependent DNA methyltransferase | 660/660(100%) | EBV0095600.1 | Type I restriction-modification enzymes |
| 11 | c14,061 – 14,558 | Toxin-Antitoxin gene | Addiction module antitoxin | 165/165 (100%) | EBV0095599.1 | Toxin-antitoxin system |
| 12 | c14,708 – 15,331 | Resolvase | Resolvase | 206/207 (99%) | WP_006811352.1 | DNA mobility |
| 13 | c15,601 – 16,365 | IS6100 | IS6-like element IS6100 family transposase | 254/254 (100%) | WP_077191101.1 | DNA mobility |
| 14 | c16,900 – 17,394 | Hypothetical protein | Hypothetical protein | 164/164 (100%) | AFI96177.1 | Unknown |
| 15 | c17,500 – 18,339 | *sul1* | Sulfonamide-resistant dihydropteroate synthase Sul1 | 279/279 (100%) | WP_000451390.1 | Resistance to sulfonamide |
| 16 | c18,333 – 18,680 | *emrE* | Ethidium bromide-methyl viologen resistance protein EmrE | 115/115 (100%) | WP_053764310.1 | Putative drug-resistant associated gene |
| 17 | c18,881 – 19,354 | *dfrA15* | Trimethoprim-resistant dihydrofolate reductase DfrA15 | 157/157 (100%) | WP_001355915.1 | Resistance to trimethoprim |
| 18 | 19,511 – 20,524 | Int-xerD | Class 1 integron integrase IntI1 | 337/337 (100%) | WP_000845048.1 | DNA mobility |
| 19 | 20,939 – 22,621 | *tniA* | TniA | 559/560 (99%) | AJD77018.1 | DNA mobility |
| 20 | 22,624 – 23,532 | *tniB* | AAA family ATPase | 302/302 (100%) | WP_000393453.1 | DNA mobility |
| 21 | 23,529 – 24,746 | *tniQ* | TniQ family protein | 405/405(100%) | WP_000801210.1 | DNA mobility |
| 22 | 24,807 – 25,421 | *tniR* | Recombinase family protein | 204/204 (100%) | WP_000904941.1 | DNA mobility |
| 23 | c25,474 – 25,710 | *merE* | Broad-spectrum mercury transporter MerE | 78/78 (100%) | WP_001087809.1 | Mercury resistance |
| 24 | c25,707 – 26,072 | *merD* | Mercury resistance co-regulator MerD | 121/121 (100%) | WP_003465059.1 | Mercury resistance |
| 25 | c26,089 – 27,735 | *merA* | Mercury (II) reductase | 548/548 (100%) | WP_092230857.1 | Mercury resistance |
| 26 | c27,732 – 27,977 | *merF* | Mercury resistance system transport protein MerF | 81/81 (100%) | WP_000654684.1 | Mercury resistance |
| 27 | c27,980 – 28,255 | *merP* | Mercury resistance system periplasmic binding protein MerP | 91/91 (100%) | WP_000735441.1 | Mercury resistance |
| 28 | c28,271 – 28,621 | *merT* | Mercuric ion transporter MerT | 116/116 (100%) | WP_001294667.1 | Mercury resistance |
| 29 | c28,693 – 29,145 | *merR* | Hg(II)-responsive transcriptional regulator | 144/144 (100%) | WP_000429838.1 | Mercury resistance |
| 30 | c29,156 – 29,605 | Hypothetical protein | Hypothetical protein | 146/149 (98%) | WP_001965932.1 | Unknown |
| 31 | c29,583 – 29,744 | Hypothetical protein | Hypothetical protein | 53/53 (100%) | EKL05168.1 | Unknown |
| 32 | c29,731 – 30,162 | Hypothetical protein | Hypothetical protein | 143/143 (100%) | WP_000834463.1 | Unknown |
| 33 | c30,159 – 31,001 | Hypothetical protein | Hypothetical protein | 279/280 (99%) | WP_001971006.1 | Unknown |

* c: indicates ORFs transcribed on the complementary strand.

### Clustal Omega Alignment of the GyrA protein of the Santiago and N16961 strains

```
GyrA_Santiago      MSDLAKEITPVNIEDELRSSYLDYAMSVIVGRALPDVRDGLKPVHRRVLFAMNVLGNDWN        60
GyrA_N16961        MSDLAKEITPVNIEDELRSSYLDYAMSVIVGRALPDVRDGLKPVHRRVLFAMNVLGNDWN        60
                   ************************************************************

GyrA_Santiago      KAYKKSARVVGDVIGKYHPHGDIAVYDTIVRMAQPFSLRYMLVDGQGNFGSIDGDSAAAM        120
GyrA_N16961        KAYKKSARVVGDVIGKYHPHGDSAVYDTIVRMAQPFSLRYMLVDGQGNFGSIDGDSAAAM        120
                   ********************** *************************************

GyrA_Santiago      RYTEVRMSKIAHELLADLDKETVDYVPNYDGTEQIPAVLPTKIPNLLINGSSGIAVGMAT        180
GyrA_N16961        RYTEVRMSKIAHELLADLDKETVDYVPNYDGTEQIPAVLPTKIPNLLINGASGIAVGMAT        180
                   **************************************************:*********

GyrA_Santiago      NIPPHNLGEVIDGCLAYIDNEAITIDELMDYIPGPDFPTAALISGRKGIIDAYKTGRGKV        240
GyrA_N16961        NIPPHNLGEVIDGCLAYIDNEAITIDELMDYIPGPDFPTAALISGRKGIIDAYKTGRGKV        240
                   ************************************************************

GyrA_Santiago      YMRSKAEIETDKNGKETIIVSEIPYQVNKARLIEKIAELVKEKKVEGISALRDESDKDGM        300
GyrA_N16961        YMRSKAEIETDKNGKETIIVSEIPYQVNKARLIEKIAELVKEKKVEGISALRDESDKDGM        300
                   ************************************************************

GyrA_Santiago      RIVIECKRDAVGEVVLNNLYANTQLQTTFGINMVALDNGQPKLFNLKEMLKCFVDHRREV        360
GyrA_N16961        RIVIECKRDAVGEVVLNNLYANTQLQTTFGINMVALDNGQPKLFNLKEMLKCFVDHRREV        360
                   ************************************************************

GyrA_Santiago      VTRRTIFELRKARERAHILEGLALALANIDEIIELIRRAATPAEAKEGLIARGWDLGNVA        420
GyrA_N16961        VTRRTIFELRKARERAHILEGLALALANIDEIIELIRRAATPAEAKEGLIARGWDLGNVA        420
                   ************************************************************

GyrA_Santiago      AMLERAGTDAARPEWLEPQYGIRDGKYFLTEQQAQAILDLRLHKLTGLEHEKILDEYKQL        480
GyrA_N16961        AMLERAGTDAARPEWLEPQYGIRDGKYFLTEQQAQAILDLRLHKLTGLEHEKILDEYKQL        480
                   ************************************************************

GyrA_Santiago      LEEIAALMHILASTERLMEVIREELEAIRAGFSDARRTEITAATHDIDMEELIAREDVVV        540
GyrA_N16961        LEEIAALMHILASTERLMEVIREELEAIRAGFSDARRTEITAATHDIDMEELIAREDVVV        540
                   ************************************************************

GyrA_Santiago      TLSHEGYVKYQLLSDYEAQRRGGKGKSATKMKEEDYIERLLVANTHDNILLFSTRGKTYR        600
GyrA_N16961        TLSHEGYVKYQLLSDYEAQRRGGKGKSATKMKEEDYIERLLVANTHDNILLFSTRGKTYR        600
                   ************************************************************

GyrA_Santiago      MKVYQLPLASRTARGKPIVNLLPLEENERITAILPVTEFSEDKFIFMATGDGTVKKTSLD        660
GyrA_N16961        MKVYQLPLASRTARGKPIVNLLPLEENERITAILPVTEFSEDKFIFMATGDGTVKKTSLD        660
                   ************************************************************

GyrA_Santiago      QFANVRANGLIALNLRDEDSLIGVDITDGESEIMLFSKFGKVVRFKESEETAVVDENGQP        720
GyrA_N16961        QFANVRANGLIALNLRDEDSLIGVDITDGESEIMLFSKFGKVVRFKESEETAVVDENGQP        720
                   ************************************************************

GyrA_Santiago      VLDENGQPEIKFKGVRPMGRTAAGVRGMKLADGDQVVSLIVPKTEGDVLTVTENGYGKRT        780
GyrA_N16961        VLDENGQPEIKFKGVRPMGRTAAGVRGMKLADGDQVVSLIVPKTEGDVLTVTENGYGKRT        780
                   ************************************************************

GyrA_Santiago      SLSEYPTKGRGTQGVVSIKVSERNGSVVGAVQVAEGDEFMMITNAGTLVRTRVAEVSQVG        840
GyrA_N16961        SLSEYPTKGRGTQGVVSIKVSERNGSVVGAVQVAEGDEFMMITNAGTLVRTRVAEVSQVG        840
                   ************************************************************

GyrA_Santiago      RNTQGVTLIRTSEGESVVGLQRIDEIEESELPEGEEALVENDAPVAQDDDGEQE    894
GyrA_N16961        RNTQGVTLIRTSEGESVVGLQRIDEIEESELPEGEEALVENDAPVAQDDDGEQE    894
                   ******************************************************
```
